# Rethinking success, integrity, and culture in research (Part 2) — A multi-actor qualitative study on problems of science

**DOI:** 10.1101/2020.02.12.945899

**Authors:** Noémie Aubert Bonn, Wim Pinxten

## Abstract

**Background:** Research misconduct and questionable research practices have been the subject of increasing attention in the past few years. But despite the rich body of research available, few empirical works provide the perspectives of non-researcher stakeholders.

**Methods:** To capture some of the forgotten voices, we conducted semi-structured interviews and focus groups with policy makers, funders, institution leaders, editors or publishers, research integrity office members, research integrity community members, laboratory technicians, researchers, research students, and former-researchers who changed career to inquire on the topics of success, integrity, and responsibilities in science. We used the Flemish biomedical landscape as a baseline to be able to grasp the views of interacting and complementary actors in a system setting.

**Results:** Given the breadth of our results, we divided our findings in a two-paper series with the current paper focusing on the problems that affect the quality and integrity of science. We first discovered that perspectives on misconduct, including the core reasons for condemning misconduct, differed between individuals and actor groups. Beyond misconduct, interviewees also identified numerous problems which affect the integrity of research. Issues related to personalities and attitudes, lack of knowledge of good practices, and research climate were mentioned. Elements that were described as essential for success (in the associate paper) were often thought to accentuate the problems of research climates by disrupting research cultures and research environments. Even though everyone agreed that current research climates need to be addressed, no one felt responsible nor capable of initiating change. Instead, respondents revealed a circle of blame and mistrust between actor groups.

**Conclusions:** Our findings resonate with recent debates, and extrapolate a few action points which might help advance the discussion. First, we must tackle how research is assessed. Second, approaches to promote better science should be revisited: not only should they directly address the impact of climates on research practices, but they should also redefine their objective to empower and support researchers rather than to capitalize on their compliance. Finally, inter-actor dialogues and shared decision making are crucial to building joint objectives for change.

**Trial registration:** osf.io/33v3m

## BACKGROUND

When performing scientific research, researchers agree to abide by principles and standards of practice. We know, however, that best practices are not always upheld (1-3). Obvious deviations from accepted practices are generally known as misconduct. But misconduct is difficult to define. At the moment, one of the most widely accepted definition of misconduct comes from the US Department of Health and Human Services 42 CFR Part 93. This definition is endorsed by the US National Institute of Health and Research Integrity Office, and defines misconduct as “fabrication, falsification, or plagiarism in proposing, performing, or reviewing research, or in reporting research results.” Nonetheless, the definition also specifies that “research misconduct does NOT include honest error or differences of opinion” (4). In other words, even in its simplest definition, misconduct remains contextual and nuanced, further complicating what constitutes research integrity. Adding to this complexity, several behaviours which cannot be characterised as manifest misconduct are also thought to deviate from research integrity. These behaviours, referred to as questionable — or detrimental — research practices, are so common in the scientific community (2, 3) that their cumulative damage is believed to surpass the damage of manifest misconduct (5). Nonetheless, questionable research practices are not univocally condemned, adding to the challenge of distinguishing acceptable from inacceptable practices.

Beyond the complexity of identifying which behaviours transgress research integrity, the causes that may lead to integrity deviations also bring confusion and disagreement. A vast body of research on the topic suggests that both personal and environmental factors are at play. Some studies condemn personal factors such as ego and personality (e.g., 6, 7-13), gender (e.g., 14, 15), and career stage (e.g., 2, 16). A few others instead believe that researchers’ lack of awareness of good practices (e.g., 17, 18, 19), inadequate leadership modelling and mentoring (20, 21), and inefficient oversight (22) are to blame. But some studies also suggest that issues embedded in the research system are at play (23). Among those, the pressure to publish (e.g., 24, 25-28), perverse incentives and conflicting interests (e.g., 29, 30-32), and competition (28) are the most frequent suspects. In light of these works, integrity seems to depend on a complex interactions between individual and social factors, climates, and awareness.

Despite the rich body of research available to explain what threatens research integrity, few empirical works target the perspectives of the stakeholders beyond researchers (33). Given the diversity of actors involved in research systems, focalising the integrity discourse on researchers inevitably overlooks essential voices.

To add some of the forgotten voices to the discourse and understand how non-researchers perceive the scientific climate, we captured the perspectives of policy makers, funders, institution leaders, editors or publishers, research integrity office members, research integrity community members, lab technicians, researchers, research students, and former-researchers who changed career on the topics of success, integrity, and responsibilities in science. We used the Flemish biomedical landscape as a baseline to be able to grasp the views of interacting and complementary actors. Given the breadth of our results, we divided our findings in a two-paper series, with the current paper focusing on the problems that affect the integrity of science (see the associated paper on success 34).

## METHODS

In the current paper, we retell the perspectives of different research actors on misconduct and on the problems which affect the integrity of science. Our data comes from interviews and focus groups with PhD students (PhD; N=6, focus group), post-doctoral researchers (PostDoc; N=5, focus group), faculty researchers (researchers; N=4, focus group), laboratory technicians (LT; N=5), researcher who changed career (RCC; N=5), members from research integrity offices (RIO; N= 4), research institution leaders (N=7), policy maker or influencers (PMI; N=4), members of the network of research on research integrity (RIN; N=3), research funders (FA; N=5), and editors or publishers (EP; N=8). The project was conducted in Flanders, Belgium, and most participants came from, or were connected with, the Flemish research system. This paper is part of a two-paper series. The full methods, materials, and participants are detailed in the associated open access paper (34).

## RESULTS

The purpose of this paper is to retell, connect, and extend on the issues that the different actors raised in our study. Aiming to maximise transparency and to minimise selective reporting, we provide numerous quotes and personal stories to illustrate our claims. The result, however, is a lengthy paper in which we explain the breadth of the concerns raised by our participants. Given the length of the resulting paper, a short summary of results is available at the end of the results section, and select findings are re-examined and extended in the discussion.

### Misconduct

#### Why misconduct matters

As we explain in the introduction, defining misconduct is challenging and very likely dependent on the context and the research culture in place. Probing directly for these complex definitions risked generating rote answers from our interviewees. Consequently, instead of asking our respondents to define misconduct, we asked them about the ‘red flags’ that indicate when researchers may be involved in inacceptable practices. By explaining these red flags, interviewees went beyond a finite list of research behaviours that they believed lacked integrity, and hinted at the reasons and personal perceptions of integrity in science.

Many interviewees started by explaining that misconduct was very difficult to detect. Some explained that the continuum of questionable research practices blurred the distinction between what may be considered misconduct, what may be punishable, and what may be acceptable despite deviating from best practices. Others sustained that misconduct had a “shifting” definition which challenged accusations of past misbehaviours. But most interviewees mentioned that the biggest challenge in detecting misconduct was the difficulty to *prove intention*. Interviewees who had to deal with cases of misconduct mentioned being able to ‘feel’ when a case was intentional, but often missed the elements to prove it. The following quote illustrate this thought.

> *But I know that there is a problem with integrity in that person. I can feel it. We have no proof. (RIL)*
>
> *We had this case once of a guy… and I, up until now, I’m still convinced that he completely fabricated his research. I know for sure that he did. But we weren’t able to prove it because it’s very difficult to prove that something is not there*. […goes on to describe the case in which all possible evidence were erased] *So there was no proof anywhere. And it was the adding up of all these coincidental things that made us believe – and in fact of course his attitude and the entire person, him as a person being was very unreliable with a lot of lies, with a lot of contradictions, stories that didn’t add up, very negatively threatening… so it was a very nasty one. And at that point you sense that there is something off. (RIO)*

Asking about red flags also allowed us to grasp what made specific practices inacceptable. We found that the reason for condemning practices ranged from general worries about the impact on science to worries about the morality and motives of the researchers. We illustrated these main positions in Figure 1, and illustrated each position with quotes in Appendix 1.

**Figure 1.**
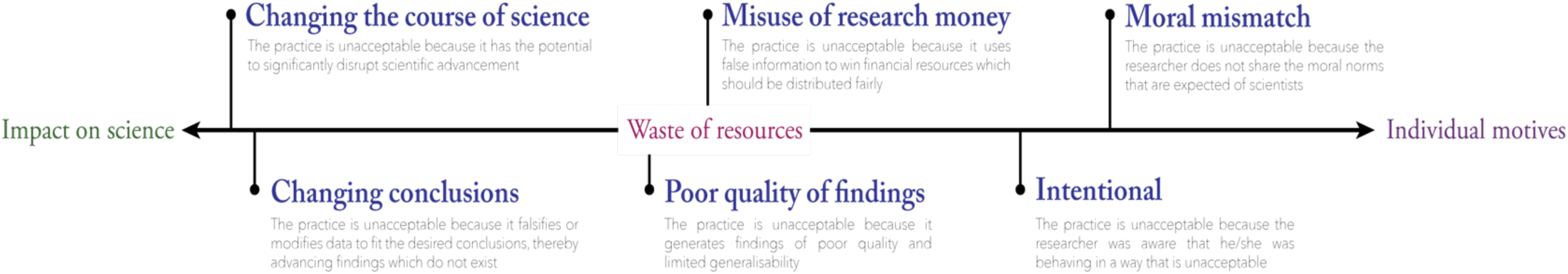
What makes questionable practices unacceptable?

The answers were mixed and diverse, but some group-specific characteristics could be observed. Among those who worried most about the impact on science, some interviewees emphasized that the potential to alter conclusions or change the course of science was what made misconduct troublesome. Editors and publishers were particularly strong on this view. Although they acknowledged the importance of intention to condemn misconduct, editors and publishers emphasized that, given their late entry in the research process, their main concern was on the effect that misconduct may have on the scientific record. This view was not only the view of editors. Some institution leaders also highlighted that not all bad intentions shape equal forms of misconduct. For example, while intentions to save efforts and be lazy can clearly harm the quality of results, they might be of a different order than the intention to change conclusions for your personal benefit.

> *“In my lab if I look, the only misconduct I’ve picked up was just stupidity. PhD students who scanned a little too short and had to go back to the scanner and thought “I could just copy-paste the bottom bit because there’s nothing on it anyway”. That’s real misconduct, but at the same time, that’s not scientific fraud. Well it was, it is scientific fraud, but he was not changing a conclusion, he was just too lazy to scan a really nice experiment [*…*] What I consider cheating is that you leave out the data that don’t suit your model. Or you make up data to get your model correctly. That is what I call cheating*.*” (RIL)*

Although both cases are unquestionably intentional, in the first case science is harmed as a side effect of pursuing a goal extrinsic to science (i.e. laziness), while in the second case science is harmed by explicitly by going against its intrinsic goals (i.e. producing inaccurate results).

The same interviewee later mentioned that small deviations, including misconduct in early stages of research, might not be so problematic since they were likely to be corrected early on and had little risk of changing the course of science (See quotes in Appendix 1). Nevertheless, research institution leaders varied greatly in their answers, and some rather sustained that intention and truthfulness of researchers were most important since they affected them as employers. Somewhere between these views, other interviewees argued that misuse of research money immediately constituted misconduct. In this regard, one policy maker or influencer sustained that any misrepresentation or duplication in an application done purposively ‘*to win money’* should be considered fraud. Another added that producing weak or low quality results which could not be generalized or used for further research was also “*sloppy or bad practice*” since the results will not represent reality. One research funder supported this perspective by adding that poor quality and delays in delivery were crucial to them since their goal was to “*guarantee the most efficient use of public money*”. Finally other interviewees focused more on the individual than on the impact on research and resources. Research integrity office members in particular tended to talk about intention or morals as the aspect that was most important to flag and determine misconduct. One interviewee explicitly mentioned that even if the conclusions were unchanged and the results were simply slightly embellished, the intention and moral mismatch was what made practices inacceptable.

#### Integrity jargon

Although research integrity office members, research integrity network members, and editors or publishers used the key terms from the integrity literature (e.g., falsification, fabrication, plagiarism (FFP), misconduct, and questionable research practices (QRP)), many interviewees, including funders, policy makers or influencers, researchers, and some institutions leaders, appeared less familiar with this jargon. They would use descriptions such as ‘changing your data’, ‘faking data’, ‘cheating’, rather than the more familiar FFP and QRP terms. Even the term ‘misconduct’ was rarely used, most often replaced by ‘fraud’. This unfamiliarity with integrity jargon may be due to Dutch-speaking nuances, or to our sampling strategy (i.e., we intentionally include interviewees who were not integrity experts in order to obtain a perspective that was unbiased by the integrity literature). Nonetheless, this finding also means that researchers working on research integrity should be aware that common terms such as FFP, misconduct, QRP, and other key terms may still be jargon to actors who are, ultimately, the intended audience.

#### What causes misconduct?

We asked our respondents *why* they think misconduct and questionable research practices happens, and whether they think it *can happen to anyone*. The main themes mentioned are illustrated in Figure 2, and illustrative quotes for each theme may be seen in Appendix 2. Since some of these themes were also mentioned as general problems of academia, we will repeat and expand on the themes later on in the ‘Problems beyond misconduct’ section.

**Figure 2.**
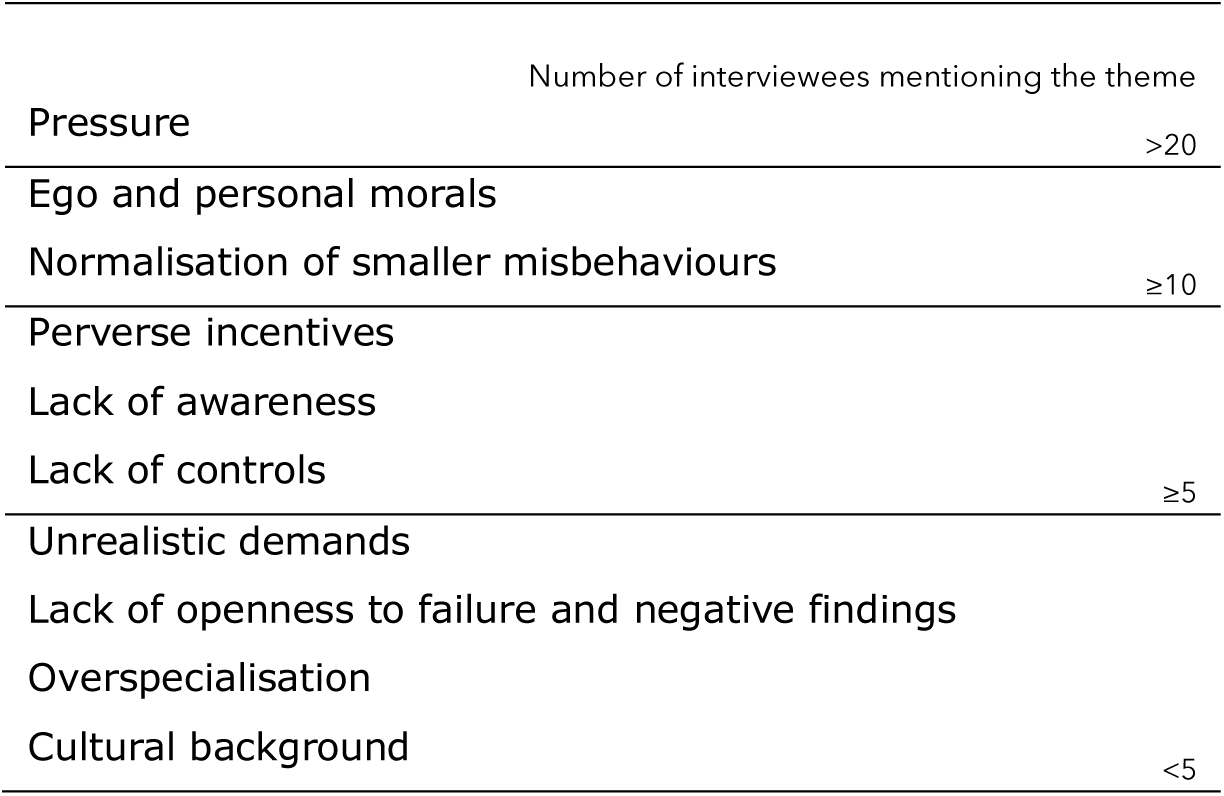
Main themes mentioned as causes for misconduct.

**Pressure** was among the most mentioned potential causes for misconduct and questionable research practices. Despite the frequent reference to pressure as being excessive and problematic, at least ten interviewees (including LT, PMI, FA, RIL, and RIO) sustained that pressure ultimately does not discharge researchers from their personal responsibilities to act with integrity. Key arguments for this position included the fact that research is not the only profession in which pressures are high, the view that pressures cannot justify moral deviance, and the perspective that even though pressures are high, pursuing research careers is a choice of which researchers are ultimately responsible. Select quotes expressing these ideas are available in Table 1. **Egos and morals**, or the ‘bad apple’ idea, was also recurrently mentioned as a possible explanation for misconduct. The high prevalence of interviewees who mentioned egos and morals might have been primed by our question ‘can misconduct could happen to anyone?’ (as we will see later), yet many interviewees spontaneously mentioned the influence of personalities and morals on misconduct. Respondents especially linked egos to ‘big misconduct cases’ such as the cases that appear in the news. **Growing tolerance of misbehaviours**, either by seeing colleagues perform bad science or by getting away with small suboptimal practices, was also often discussed as a catalyst for detrimental practices and misconduct.

**Table 1.**
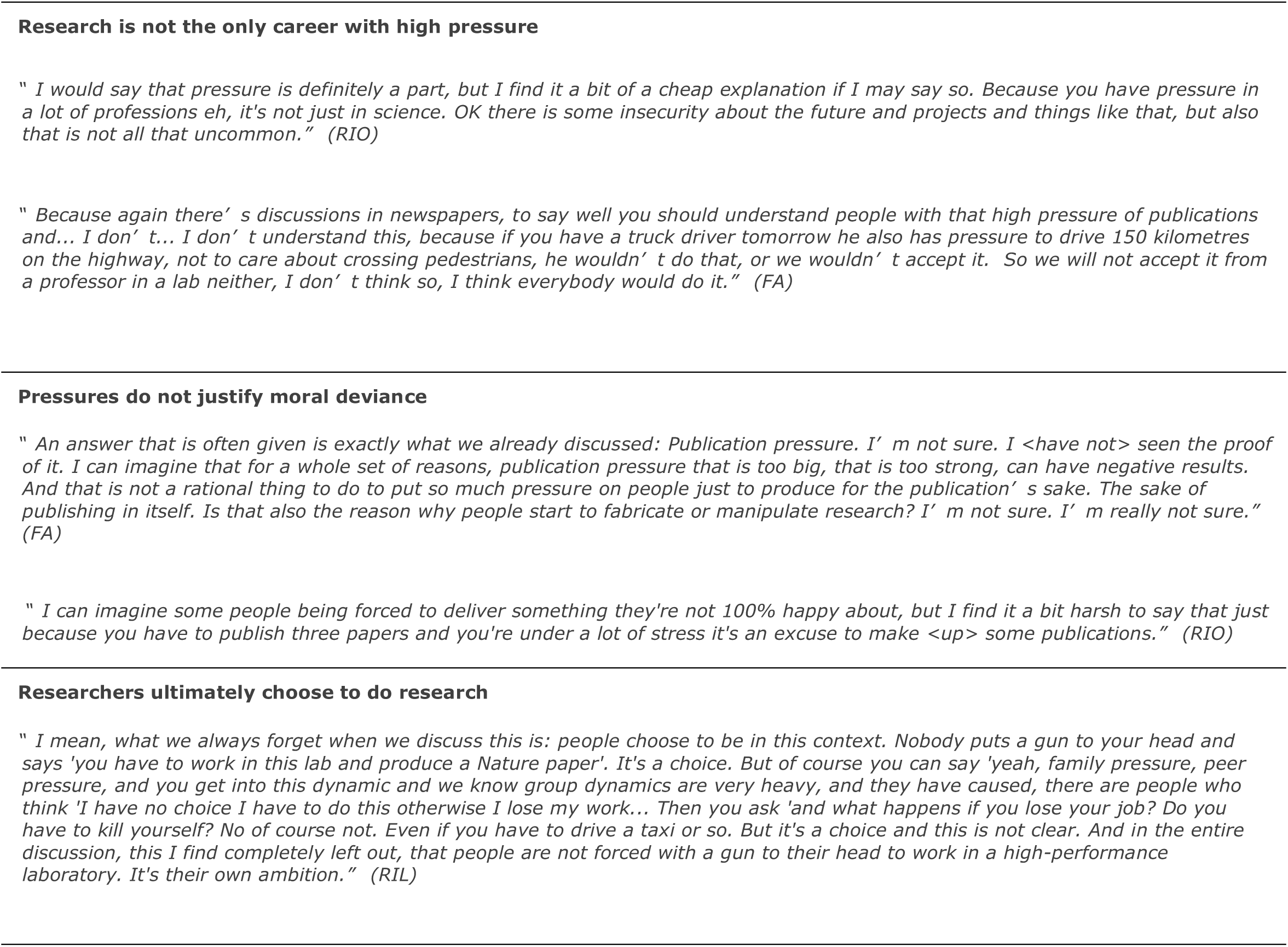
Select quotes supporting that pressure may not be an excuse for misconduct

Beyond these three major culprits, additional determinants of misconduct were raised. Among those, the perverse effects that unsuitable incentives may incur, added to a lack of control for compliance, were thought to make misconduct a low-risk high-gain prospect. The overspecialization of research fields was also thought to challenge monitoring and reproducibility. Finally, research culture were also thought to threaten integrity, for instance through the cultural background of researchers or students, the lack of openness for failure and negative findings, and the lack of realism in expectations and demands. We will revisit these three themes later on since they were also mentioned as more general problems of science.

#### Is everyone subject to misconduct?

When asking respondents whether misconduct could happen to anyone, the answers were varied and contradicting. Although respondents identified pressures and problems from the research culture and environment as major causes for misconduct (cf. supra), most respondents also supported that certain types of personalities were more prone to misconduct than others. The extent of this perception, however, varied from interviewee to interviewee, and appeared to be linked to personal experiences with misconduct cases rather than to actor groups.

First, some interviewees perceived researchers as inherently good by default. Statements along the lines of “I believe in the goodness of researchers” (RIO), “She was a real scientist… I could not believe that she would ever, yeah, <commit> misconduct on purpose.” (RCC), or “I find it very hard to believe that somebody who would go into science, go into research to intend really to go and do wrong things.” (RIO) illustrate this perspective. Nonetheless, the same interviewees later explained that despite researchers’ inherent goodness, academia sometimes placed so much pressure on researchers that it may push them to deviate from integrity. Corroborating the view that research culture and environments may drive virtually anyone to commit misconduct, other interviewees were more explicit in linking propensity for misconduct to individuals. These respondents sustained that, although pressures still played a key role in misconduct, certain researchers were more prone to misconduct than others.

> *“I definitely think there is a pathological end of the spectrum*.*[…] But I also think that there is so much pressure, especially on people at the beginning of their careers, that I don’t think anyone is completely immune to actually committing something” (EP)*
>
> *“The truly white and the truly blacks are rare. […] many people will be willing to cut a small corner somewhere in an experiment. But really cutting a corner meaning ‘I come up with an answer that I don’t have yet, but I assume it will be this and I’ll give myself the data for free’*… *I think requires a mentality”. (RIL)*
>
> Finally, a minority of interviewees believed that individual characteristics were the biggest (if not sole) determinant of integrity. Although this perspective was only supported by a few interviewees, it suggests that integrity is sometimes perceived as independent from training and climates. Supporters of this view questioned the benefit of training and support in promoting research integrity, and rather sustained that, to build good researchers, institutions must choose the right individuals.
>
> *“Sloppy science, first and foremost is the product of sloppy scientists. It’s not the product of a system, it’s the product of a person. […] there are persons who are striving for high levels of integrity, and there are people who are not doing so” (PMI)*.
>
> *“Integrity is in the person. […] integrity is something that is in you. You have it or you don’t have it. I mean you have it, it’s there. And when you don’t have it, you don’t have it. So we cannot create integrity, it’s something that’s in the people. Working together and being involved, that’s something [universities] can create by offering a structure. But I’m a strong believer that the integrity is inborn, it’s in you”. (RIL)*

Discussions on misconduct sometimes appear to be obvious and consensual. Yet, our interviews revealed that perspectives, knowledge, and convictions about misconduct can vary greatly between individuals and actor groups. Not only are the terms used to talk about misconduct still jargon to many research actors, but the views on why misconduct matters, what causes it, and who is susceptible to it also varied greatly between interviewees.

### The problems beyond misconduct

In performing our interviews, we noticed that what respondents were most concerned with was not ‘misconduct’ per-se, but rather a number of more general problems that affect the integrity of research.

> *“The experience that I have in research is that really [misconduct] is exceptional. It makes… It’s breaking news, because it’s something that we, in the community of research, we consider inacceptable, but it’s rare. It’s rare*.*” (FA)*
>
> *“The very serious misconduct is not such a big problem. It’s… it’s more the grey area that is a problem because of, yeah, the amount of… of, yeah, the bad practices*.*” (RIO)*

Indeed, respondents discussed what they found problematic in research and what frustrated them much more spontaneously than genuine misconduct. They expressed these problems and frustrations throughout the interview, even during discussions on success and responsibilities, often forcing the interviewer to re-focus the discussions on the ‘positive side’ of science. Results from the following sections are thus based on spontaneous reflections expressed throughout the interview rather than limited to specific interview questions.

#### A tight connection between success indicators and threats to integrity

The first thing we noticed when analyzing the problems and frustrations raised by our respondents is that many of them are intimately connected to the way success is attributed in science. In Figure 3, we linked the different themes of success which we reported in Part 1 of this study (34) to the different problems or frustrations mentioned by interviewees. Despite the oversimplification of this illustration, we can see at first glance that the two topics are highly interconnected. In fact, not only are determinants of success seen as aggravating some of the problems mentioned by interviewees, but some of the problems mentioned are also seen as blocking success in science. In addition, problems appeared to influence one another, escalating into bigger issues until some of them became big enough to generate misconduct. In this regard, some of the problems described as causes for misconduct were first described as general problems of science.

**Figure 3.**
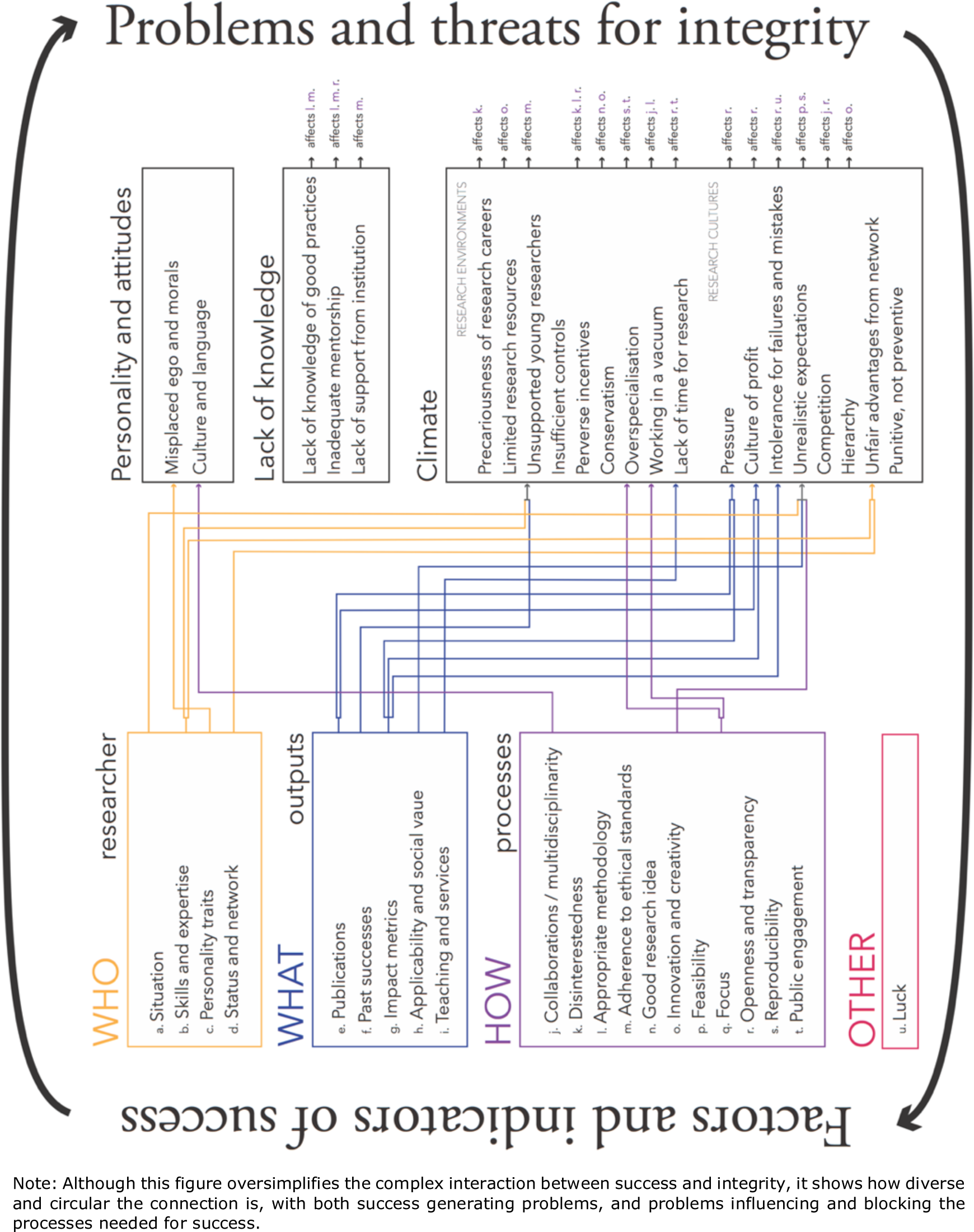
Simple depiction of the complex interaction between the factors and indicators that currently define success and the problems which currently threaten research integrity.

#### Three categories of threats to integrity in science

Interviewees raised a number of problems which could potentially threaten the integrity of science. We organized the different problems mentioned in three big categories: problems related the *personality and attitudes*, problems related to *a lack of knowledge from researchers*, and problems related to *research climate*s, which include research environments and research cultures.

##### Issues related to the personality and attitudes

Interviewees mentioned a number of individual characteristics which could be problematic and might impede on the integrity of research. We have already mentioned a number of these issues — such as misplaced ego and morals — above when discussing individual propensity to misconduct. But a few points raised conflicting dualities with what was believed essential to success. For instance, interviewees supported that ambition, passion, and tenacity were key elements of success (34). Nonetheless, they also supported that hyper-ambition or excessive desire to be successful could bias conclusions and encourage researchers to loosen their integrity.

> *“That’s very important, because we’re always talking about misconduct as if it’s deliberate, as if you’re cheating, I think maybe the most dangerous thing in research is in your wishful thinking, of self-fulling prophecies, you want it so badly that you will see it, you will see it the results, if you’re out in a complex…” (FA)*
>
> *“And the good researchers, like I said, they’re really passionate, they only think about their own research, they want to get things done, they want to get their results, so… What we usually see is that people then don’t really follow the rules as they should, so they don’t see why these rules are important. […] It’s always that this researcher is really heart to… motivated to get the results done, and then bypasses procedures and rules, and that they don’t see why these rules are in place and why they’re so important to have them, why it’s also protecting other people in the field… So this is mainly I think one of the reasons…” (RIL)*

Some respondents also associated attitudes with particular cultural backgrounds. For instance, some interviewees proposed that the perceived seriousness of detrimental research practices may differ between cultures.

> *“…Sometimes I find that it’s a matter of some cultural differences, in some cultures it seems that every meaning is justifiable to achieve the goal, and so they are trying to do anything they can do just to get their research published. So they will falsify data without a problem [laughs], they will not hesitate…” (EP)*
>
> *“I think [certain cultures] have this mentality that it’s almost, you honor somebody by plagiarising them. And they just want to get their diploma so they can do a post doc in America*.*” (RIL)*
>
> *“I dare to say that [different cultures] have a slightly different opinion about rules*.*” (RIL)*

Cultural and language differences were also mentioned as challenging the ability to communicate and increasing the risk of loneliness, misunderstandings, and mistrust. For instance, laboratory technicians mentioned that cultural and language differences could decrease students’ willingness to ask questions and disclose mistakes, thereby increasing the cumulative severity of mistakes and the temptation to conceal them. But despite raising these personal issues as a potential risks for integrity, interviewees failed to propose concrete solutions or improvements for minimising the risks resulting from personalities and attitudes.

##### Issues related to a lack of knowledge

###### Lack of knowledge of good practices

Several respondents mentioned that researchers were sometimes unaware of good practices. Lack of knowledge of good practices was not referred to as a problem of the individual who lacks insight in his/her own behaviour, but as a systemic issue caused by insufficient training and inadequate mentoring within the larger scientific community.

###### Insufficient support, mentorship, and guidance

Most concerns related to the lack of knowledge of good practices pointed to a lack of mentorship and guidance for early career researchers. This issue was discussed on different levels. One the one hand, students mentioned that they lacked guidance, support, and time from their supervisors. PhD students and researchers who changed career were especially vocal on this point.

> *“Well it was generally just my supervisor messing up. That was just the worst. Always the worst. Always. (laughs) And I’m not telling you… you know. So, not responding to emails, you know, for a very long time. Not being present. Not giving any useful feedback, if they give feedback, giving feedback that just makes your work worse instead of better… Not knowing how to supervise basically*.*” (RCC)*
>
> *“I think everything I learnt, I learnt because of doing myself. I expected when I started my PhD project, that I would learn a lot from my supervisors, but now at the end of my PhD I think I didn’t really learn a lot from them, so I’m a bit disappointed about that*.*”. (PhD)*

Although the lack of mentoring in such cases is not necessarily causing an unfamiliarity with good practices, young researchers often felt lonely, stressed, and frustrated about the lack of support they receive. Loneliness, in turn, was described as a possible red flag that should be considered immediately.

> *“I get suspicious when PhD students are complaining, for instance, or feel alone or feel pressured to do things. Of course, in a certain sense this always happens. If you ask my PhD students, they will also say there are moments in which they were alone or pressured, so you cannot really prevent all of that but if that becomes too big, then I think there is something wrong” (RIN)*

On the other hand, researchers themselves mentioned that they lacked support and guidance on how they should meet integrity and ethical requirements. For example, one researcher mentioned that funders increased the number of “tick boxes” without increasing training and capacity.

> *“Just having you as a researcher filling all these tick boxes, and not being responsible*… *[Funders] really should work on that. Also the same goes now for the data protection. They will make an extra box, and we should think that everything is arranged for data protection while no university in Flanders is ready for that by May 28th*.*” (Researcher)*
>
> Along the same line, research integrity officer members worried that integrity standards were too disconnected from core research training. They sustained that integrity training generally come in the form of specialized courses, when in fact they should be integrally embedded throughout the research training in order to become “*part of the research process*” for every researcher.

##### Issues related to the research climate: Research environment

The third broad category of problems raised by our interviewees were linked to the research climate, targeting both the working environment and the research culture.

###### Precariousness of research careers and limited research resources

The **precariousness of research careers** and the constant insecurity linked to short-term contracts and scarce opportunities for advancement was a recurrent issue mentioned by our interviewees. Policy makers and research institution leaders were particularly concerned about this issue. One policy maker explained that, in Flanders, the number of students completing a PhD highly exceeds the number of academic positions available, even though the current number of PhD students in Flanders is below the recommendations of the Organisation for Economic Co-operation and Development (OECD). Another policy maker added that the ambition to continue in academia is the default option for PhD students. He further explained that this is problematic since there are “*phantom pains attached to it. People think that it’s kind of a lost battle — a defeat — when they leave university and go to work in a company, or go to work for another agency*.” Trying to find solutions to the problem, some interviewees supported that the lack of opportunities in research may result from **limited research resources**, and that investing more money in research would help solve the problem. Nonetheless, other respondents concluded that because of the way the system is organized, capital investment in research would not necessarily solve this problem.

> *“Because research is a human activity, more money into research means also more people getting involved in research. I’m always amazed to see that people think that this should lead to more academic careers in research. That’s a kind of incongruence because of the fact that the public sector cannot bloat universities, in the sense that we cannot multiply by an order of magnitude the amount of positions available*.*” (PMI)*

###### Unsupported young researchers

Adding to the lack of stability and security embedded in research careers, the **struggles of early careers** was another important theme in our interviews. Young researchers and former-researchers mentioned that they felt unsupported while juggling with too many tasks to be able to focus on the outputs required for advancing their career. Early career researchers, former-researchers, and post-doctoral researchers also worried that their modest output records disadvantaged them in the fierce competition needed to secure grants and careers in academia. As a result, on top of the duties of early adult life (building a family, buying a house, caring for aging parents, etc.), young researchers struggled with an insecure future, excessive pressures for output, insufficient resources, and the inability to compete with established researchers.

> *“There’s a certain starters package that I got, but it’s not enough, you have to find your own money which is very difficult because you don’t have the publication list. […] Because the first thing [funders] do is, they look at who is asking, and then at your resume and then they say oh no too junior or not well enough established in the filed or, you know, stuff like that. […] you need money to publish, you need to publish to get money, you know, it’s a circle*.*” (RCC)*
>
> *“Yeah so for me it’s because I’m in this end stage, the insecurity of the future is really something that I’m struggling with. Not every day, or not all day every day, but every day at least 5 minutes (laughs). […] The fact that you don’t have a permanent position is also really ambiguous about it. I would like to have a little bit more future, and also not to have to find my own money all the time because I have the feeling that I’m not actually doing something myself. I’m constantly finding and looking for more money, so to hire people who are actually doing something*.*” (PostDoc)*

Most former-researchers said that the desire for a stable career with a sane work–life balance influenced their decision to leave academia.

> *“Why should I stay in the academic world, why should I go?. […] If I go for the academic world, I’m going to have to tell my wife, that was pregnant [at the time…], I have to tell my wife “well we’re going to a financially uncertain situation for at least 10 to 15 years. And maybe when I’m 30 or 35 and I have said no to you an enormous amount of times, I’m going to be so successful that I can say ‘It’s ok now, we can pay the bills*.*’ But I’m still going to say no to you because I have to compete with the other people. Whereas if I choose another life or career, you get, for example a contract that lasts for your entire life, and you can build your life. You can start building your life. You can settle in a way, you can… You can make plans. Whereas in the academic world you can only make plans for 2 or 3 years. And that was the kind of life that I didn’t want to <live>*.*” (RCC)*

Two of the interviewed former-researchers admitted having been mentally and emotionally affected with symptoms of burnout, and all recalled a certain distress from their time in academia (we will get back to the emotional distress when discussing unrealistic expectations below). One interviewee proposed that this distress may be accentuated by “*the enormous discrepancy we have today between the job security of professors and the job non-security […] of PhD students <and> post docs*”. This interviewee recalled stories of supervisors who continuously reminded their students that they could be replaced anytime if they didn’t meet expectations. Such security discrepancy was also thought to create an environment in which young researchers may not feel safe enough to be open or transparent about issues and mistakes — a problem we will target later when discussing the research culture. Finally, there appeared to be strong emotional implications for researchers who decided to quit academia. Even though all former-researchers interviewed expressed a great sense of relief from leaving academia, most sustained that the decision to leave had been difficult to make since it would be perceived as a failure in their career. The emotional involvement was often linked to a sense of personal disappointment or shame, rather than to a frustration against an unrealistic and unsupportive system.

> *“I am the idiot that gave up [a professorship]. That’s what it is, I worked my entire career to get at that point, I was in it for [a few] years and I gave it up. And so many people in the academy want to be in that position, and I gave it up. What kind of an idiot am I?* […later in the discussion…] *In the end I was like […] What’s the chance that I’ll ever help any patients, because that’s basically why we all start doing it, to make a difference. But that’s for the happy few, and those happy few have big names behind them and get money. They are not struggling to be at home, to put children to bed or whatever. The daily things that were too hard for me, and now that I don’t have to do it anymore I’m a happier person. So maybe I’m not a real, real scientist*.*” (RCC)*

In this last quote, the perception of not being a ‘real, real scientist’ clearly expresses that researchers who leave academia tend to blame themselves rather than the system’s unreasonable demands, a perspective which further deepens the wound and pain from leaving. In sum, the strength of the aspirations that young researchers hold to continue in academia may increase their vulnerability by imposing escalating expectations upon themselves. Knowing that less than 10% of PhD students will be able to pursue academic careers, the current dynamics clearly generate disappointments, self-doubt, and emotional distress among early career researchers for whom the future is uncertain.

###### Inefficient controls and perverse incentives

Issues around **inefficient controls** were also raised by a few stakeholders who sustained that misconduct and detrimental practices often go unnoticed or unsanctioned. Research integrity officers complemented this idea by mentioning there are also insufficient incentives for integrity.

In fact, **current incentives** were often thought to discourage integrity. One interviewee in the editor and publisher group mentioned that, at the moment, researchers are incentivized to find “*big bold claims*” and to publish in “*very selective journals*”, which led to a number of low quality research practices such as performing research on smaller populations or choosing inappropriate statistical controls and analyses to inflate significance. We will get back on this point later on when discussing issues of unrealistic expectations and the culture of publish-or-perish that results from such expectations. Along the same lines, an institution leader mentioned that “*short term financing situations*” which expect high publication outcomes may be the “*worst perverse incentive you can give a scientist*”.

###### Conservatism

Adding to the above concern, the core functioning of funding distribution was criticized for being conservative and discouraging high risk research (i.e., research with important possible outcomes but with high potential for yielding negative results). Interviewees considered that high risk research was important for scientific advancement and innovation, but they worried the reliance on experts for reviews decreased the chances of obtaining funding for high risk or simply unusual research.

> *“…peer review has this tendency to be a little bit conservative. Because since you have experts in your panels, people who already have proven themselves […] and also mostly are senior people, they can also sometimes, not all of them because you shouldn’t generalize, but sometimes they can get a little bit conservative. Because they think that they have found the holy grail*.*” (FA)*

One policy maker proposed that the problem also came from within institutions, supporting that “more and more, institutions, universities don’t want to fund high risk research. So they only want to fund research that gives good results that can be used for society and so on.” As a coping strategy, both PhD students and researchers admitted having heard of situations where applicants “get funding for something that’s already proven, and they just explain it and they turn it… they describe it in such a way that it’s new, and then they get funding” (PhD student), or where researchers “write a project where <they> already have the data and… so <they’re> asking money for something that <they have> already done.” (Researcher). Researchers considered that such coping strategies were problematic because they limited innovation and prevented new research groups from obtaining funding in topics that were investigated by other, more established groups.

> *“You need to show that you have every technology in hand to do this new idea. And this is really a problem for me. I think that many researchers are now playing at the safe side. Because they already have shown that they work in this field, they will continue on this field, and they will not go broader, because probably they will not get funding because it’s a new idea and they don’t have any evidence at work*.*” (Researcher)*

In response to these concerns, one research funder stated that there also existed private, smaller funders which “*could, and therefore also probably should <be> somewhat more risk taking than public funders*.”

###### Overspecialisation, working in a vacuum, and lack of time for research

Interviewees also shed light on a problematic interplay between overspecialisation, isolation, and lack of time for research. As introduced in the potential causes for misconduct, **overspecialization** was criticized for potentially deterring the replicability of research, thereby undermining the detection of mistakes and misconduct. But overspecialisation was also criticized for increasingly isolating researchers from one another and discouraging collaborations. Interviewees often felt that researchers **work in a vacuum** rather than within a shared community. Evidencing this idea, PhD students supported that research “*is sometimes a bit lonely*” and that they were often unaware of the research that was happening around them. An interviewee from the RIO sustained that “*very often […] researchers don’t even know what is happening within their own buildings*” and that this isolation probably lead to unnecessary duplication and waste of research resource. Working alone also means that researchers are expected to have highly versatile abilities to be able to coordinate and respond to the expectations of their position.

> *“The advantage of academics is that you have many tasks, but this is also a disadvantage. Sometimes you have to do everything, you have to be good <in> English and <grammar>, in statistics, in everything, and*… *which is not always our expertise and also neither our interest*.*” (Researcher)*

Another researcher added that the lack of collaboration also reduced the possibility of blinding experimenters and increased the risks of bias. But beyond research multitasking, the three pillars required to be employed in a university — i.e., ‘Teaching’, ‘Research’, and ‘Services’ — also sparked the debate.

> *“The problem is that today we cannot deliberate between those three pillars. And I believe that if you’re excellent in education and you spend 80% of your time in education and you do only 20% in research, and you don’t get your criteria for research but you overdo your criteria in education, why not make a balance?”. (RIL)*

This lack of flexibility played an important role in the decision to quit research of one of our interviewee. Asked to teach approximately 80% of the time, this interviewee sustained that there was too little time left for fulfilling research requirements. This interviewee further sustained that the lack of flexibility from the three pillars of research careers (i.e., everyone is expected to perform research, teach, and contribute to services) neglected personal skills and preferences.

> *“I believe there are very good researchers that have to teach and that suck at it, and I believe that there are good teachers that have to do research and suck at it. Or at least are not top notch at it. But no we all have to be equally good at both and we all have to divide our time exactly the same. […] I struggled at doing everything the way I want to do it. I want to do everything in a good way. And when you have to do teaching and research I didn’t manage. I didn’t manage to do both in a good way*.*” (RCC)*

Finally, with such diverse tasks and pillars of expertise, and with the bureaucratic demands of research work, researchers felt that they **lacked time to actually do research**.

> *“I think the research part you’re so passionate about it and then, you know, you feel, you always have to fight to get your time to do it. And there is many, so many things that always come unexpectedly, or expectedly in between, that disable you from writing that article, or from doing your field work yourself*… *[…] because we have to do also education and we have to do managing tasks, and then we have a curriculum reform, then we have to think also about the new education, and then we have, we are responsible for clinical placement and things go wrong on the clinical placement, and then*… *I mean because I’m juggling many balls, it always*… *seems like I always, for one reason or the other, have to be juggling those balls instead of being able to do, to spend more time on my research. And we have a tremendous amount of meetings*… *The amount or time that I’m sitting in the meeting room is … (sighs)” (Researcher)*

Ultimately, this lack of time played back and aggravated a number of issues we just mentioned, such as inadequate mentorship and the difficulty to build one’s status as an early career researcher.

##### Issues related to the research climate: Research culture

Deeper into the habits and customs of researchers, several issues embedded in the culture of research were also seen as problematic. Once again, Figure 3 showcases a few of the interactions between issues embedded in scientific culture and current indicators of success.

To begin, pressure to perform — and especially to deliver — was mentioned by almost all participants as creating a general publish-or-perish culture. Excessive pressure was further described as creating an atmosphere where profit and positive results are expected, and where failures and mistakes are not tolerated. We will get back to each of these points later on.

###### Pressure

**Pressure** and the culture of publish or perish were the issues that were mentioned by the biggest number of interviewees. Interviewees described such pressure as potentially causing misconduct, as threatening the quality of science, and as impeding on researchers’ health and happiness. These ideas have often been targeted in past literature on research misconduct and integrity. Our study, however, provides a new perspective on pressures which differs from the traditional views present in the literature. By listening to multiple research actors, we discovered that pressures are multilevel and that they affect more than researchers alone. In fact, the publish-or-perish culture fuels a cascade of expectations and demands which increase pressures on a broad range of research actors. Starting with students and researchers, we first found that pressures did not only come from the institutions and superiors, but sometimes came from the researchers themselves, in the form of personal aspirations and ambitions. On a second level, students and researchers expressed feeling substantial pressure coming from the supervisor and the institution. But institution leaders also expressed that they felt, as an institute, a pressure to deliver more and faster in order to promote the excellence of their research and their attractiveness to the international research community. One research funder explained that in Flanders, where institutional funding depends largely on research outputs^1^, institutions must continuously increase their outputs in order to keep their share of structural funding.

> *“…to even conserve [their] share, make sure that [they] will not get less than the previous year, [universities] have to work always harder, [they] have to produce more publications. Because if [their] competitors — other universities — produces more than [them], then the share of that same amount of money will decrease” (FA)*

One level further, funders expressed that even they felt pressures. The increasing number of applications for funding increased workload and generated internal pressure and struggle to find adequate peer-reviewers. Journals expressed a similar concern, stating that the pressure to publish and the current focus on quantity often led them to receive more manuscripts than they could review, and forced them to use greater scrutiny to ensure the quality of their publications, but also to charge higher author processing charges and subscriptions. This whole circle of pressures then appears to link back to policy makers. Specific to Flanders, policy makers were criticized for the distribution key they use to distribute funding between universities (the BOF-key, which stands for special research funds). This distribution key was said to be *“the reason that publications are so paramount in the assessments”* (PMI). Nevertheless, policy makers and influencers clearly expressed that although they were aware of the criticism generated by the BOF-key (and were currently working on a revision^2^), it was its inadequate usage (i.e., using its parameters at the individual level) that was at the source of the problem.

> *“The BOF-key was actually only created to divide the money under the universities. And what we see is that the same parameters are being used within the universities themselves to fund the individual researchers. That was never our intention… So that’s the negative effect of this key that we never wanted*.*” (PMI)*
>
> *“The BOF key is just one thing. It’s a distribution rule that has to divide a pot of money among five universities. In one way or another, you will always need some distribution mechanism. The BOF Key — That’s also why we never report on individual researchers — but what the BOF key does is just aggregated at the level of a university: count PhD output, count publication output (certain type of publication output I’ll come back to that). […] I know from hearing and feedback that I get that certain institutions try to, what I would call extrapolate, or interpolate the BOF Key into individual level research output. I think that’s wrong, that’s even stupid. But it may happen. But the BOF key it’s not there to do this*.*” (Another PMI)*

Consequently, even though the pressures and the culture of publish-or-perish were raised by nearly all interviewees, the root of the problem appears to be transferred from one actor to the next. This circle of blame further seemed to create a mis-appreciation of individual responsibilities and actionable solutions, leaving most actors feeling frustrated helpless. We will discuss this problem further later on.

###### Culture of profit

In tight connection with the pressures and the culture of publish or perish, the whole **culture of profit** that characterizes current academia was also questioned by a few interviewees. More specifically, the emphasis on profit and outcomes was seen as potentially undermining the care and consideration that should be given to researchers themselves.

> *“In research you’re not making research results, you’re making good researchers. And you have to develop and support the people, and not just the research. And I think that that entire culture of care is missing too much. We see them too much as producers of research results, instead of ‘we are making a good researcher that will, hopefully go on a lifetime making good research results’*.*” (RIO)*

This forgotten need for care easily links back to the lack of support faced by young researchers, the precariousness of research careers, and the lack of support for meeting integrity requirements, while it also links forward to another problem: the lack of tolerance for failure and mistakes.

###### Intolerance for failures and mistakes

Interviewees from all actor groups spontaneously explained that **failure, negative findings, and mistakes are almost invisible in science**. Yet interviewees also sustained that failures were *very “important*”, “*valuable*”, and “*interesting*”, and that they could act as “*a motor to drive you to success*” (PMI). Lack of tolerance for failure and mistakes was even thought to be a potential incentive for falsification of results (see Appendix 1). One researcher told of a personal experience when discovering a mistake in a team project. From the story, the different reaction and the overall worry that mistakes can generate in science is obvious.

> *“I think what is also a problem is the fact that it’s still a ‘taboo’ I would say, just to come open with the fact within research “I made a mistake” in the past. We had something in the past in our group that there was… suddenly there was… everybody thought that a measurement was wrong. Something in the system and all of the data that were captured were therefore wrong, and <these> were already data which were published. And then it should be decided what to do. Should we do a correction to the journal or not? And there was a lot of pressure from the professors, because it came higher and higher in the university, and some people were afraid, and some were like ‘Whatever!’, and everybody had another opinion, the PhD students just had to follow*… *But I feel the big difference between some people who were very ethically committed, like we have to correct it and we have to send it to the journal, and others were like ‘nobody will see it, it’s in the past’, and*… *Yeah, I saw a lot of things which should not have been happening*.*” (Researcher)*

Although the fear of mistakes seemed deeply engrained in the research culture, most interviewees sustained that efforts must be made to normalise mistakes in science. One former-researcher eloquently summarised this idea by stating that “*If one place in our world should be a place where people are free to make mistakes, even though we pay them a lot, and we hope they don’t make mistakes, then it’s [academia]*.” (RCC), further advancing that intolerance for failure was “*not justifiable*” in academia. The **under-appreciation for negative results** was also mentioned very frequently. Resonating current views from the literature, interviewees worried that unpublished negative results wasted research resources and potentially endangering research participants. Yet, on the research floor, the apprehension for negative results was still palpable. Researchers, research students, and lab technicians described negative results as highly frustrating or as ‘*unlucky*’ (see the discussion on luck in 34) and admitted that projects with negative results were often abandoned early. The quest for positive results also influenced research designs. Students admitted with unease that many experiments seemed designed to ensure publications rather than scientific relevance. Researchers and lab technicians added that data fishing and selective publication were common practice, even sometimes part of the strategies required for success. But when asked about responsibility, interviewees once again seemed to pass the ball to one another. Researchers claimed that they were pushed to look for positive findings since journals would not accept negative results and funders expected their projects to yield positive findings. Nonetheless, both journals and funders refuted this perspective, rather sustaining that they would not mind about the results as long as the experiment was performed well. Editors and publishers added that they rarely, if ever, received manuscripts with only negative results. One interviewee even told the story of a new journal dedicated to negative results which had to be shut down because it received “*no submission whatsoever*” (EP). The issue thus appear to be deeply embedded within the research culture, possibly even budding at the micro level within the research teams themselves.

> *“If you’re really interested in the success of research environment, it’s an environment that says ‘you don’t have to be successful’. ‘You may fail. And it’s OK. As long as your research methodology is accurate’. […] Now you see that the rector for example is also saying this, so I think change is coming in a certain way… In a certain way. But I’m not that sure if it’s really coming because the culture is defined by your promoter. […] You may get trainings every single day. If your promoter or the head of the lab doesn’t agree, then it won’t happen*.*” (RCC)*

###### Unrealistic expectations

Intolerance for failure might be a simplistic expression of a bigger problem: science builds **unrealistic expectations**. In fact, interviewees mentioned that too much was expected from researchers, potentially leading to integrity deviations, frustrations, or even burn out. Different forms of expectations were perceived as being excessive and unrealistic. First, **expectations of high yielding results, and extraordinary findings** were considered to be embedded in the core of how science is evaluated.

> *“Researchers are incentivized to really get something that is extraordinary, and ground-breaking. And let’s face it, all the research in biomedical research is not ground-breaking and extraordinary. Most of it is not*.*”(EP)*

Second, **expectations that researchers should work out of passion without personal benefits** also surged from our interviews. For example, an institution leader mentioned that institutions “*need people with commitment who <participate in services> for the honour*” without expecting personal gains or compensation. A policy maker added that he “*would never call ‘doing science’ a job* […] *being passionate about science is almost like being an artist. You live in poverty because you want to pursue your art*.” In other words, research was not seen as a regular career but rather as one that is built on devotion and personal sacrifice for the greater benefit of science. Many interviewees expressed the expectation that researchers work outside ordinary schedules^3^, travel abroad regularly, and eventually even rethink their work–life balance.

> *“I think people have to realize when you do a PhD, it’s a stressful thing, you really are going to get the highest degree there is at a university, it doesn’t fit between 9 and 5*.*” (RIL)*
>
> *“I’m somewhat older, but I have the impression that younger people have […] somewhat a different work/private balance than I had. And I think that people sometimes could put more energy in their work*.*” (RIL)*

Unfortunately, such expectations of personal sacrifices were not benign on researchers and research students. Researchers and students explained that it was difficult to conjugate their professional and personal life, and that they sometimes felt the need to sacrifice the latter to ensure their professional survival. As we have briefly discussed above, three researchers who changed career mentioned that the difficulty to keep a sane work–life balance played a significant role in their decision to move away from academia, with some adding that such excessive demands affected their well-being.

> *“I was stressed out completely […] I went to the doctor, [I] was on antidepressants, [I] was in therapy…” (RCC)*
>
> *“I got a therapist and I worked through it with her and, you know she said… Like whenever she said ‘Maybe you want to start thinking about [your work]?’ I would just start to cry, so she was like OK, too early! […] Yeah. It was awful*.*” (Another RCC)*

But even those who suffered the effects of excessive expectations tended to perceive “real scientists” as those who should give more than they could. Worrying about this unrealistic perspective and about the implications that unattractive research careers may cause in the long run, one policy maker or influencer advised that researchers should be given equivalent benefits than other individuals on the job market.

> *“Why would I choose to do, to start a career in an area that positions are limited, promotions are limited, high positions are limited, and it’s precarious. I have to bring funding, I have to get contracts*… *So*… *it’s not only the lack of interest — if it is there — from the younger generations, it is also what is the tomorrow. And this is part also, I think, of a scientific governance and a scientific culture issue. Those*… *We should not consider that researchers are somewhat a different part of the population or that they are saints, that they will sacrifice their wellbeing and their participation in the pleasures of a good economy just because they love science. I think this is very naive*.*” (PMI)*
>
> While it is beyond our purpose to determine whether researchers should, or should not sacrifice their personal life for their career, our interviews show that this expectation is still alive and that it affects researchers’ well-being.

###### Competition, hierarchy, and advantaged due to networking

A few additional problems were linked to the social relationships which characterize the academic culture, such as **competition, hierarchy**, and **advantages due to networking**.

The issue of **competition** raised mixed reactions from our participants. On the one hand, some interviewees mentioned that competition was a necessary element of academia as it drove productivity and excellence while imposing limits on the authority of single researchers. But on the other hand, competition also challenged researchers’ openness. For one former-researcher, competition was the determining factor for leave academia.

> *“Competition in the academic world is so strong, so fierce that in the end I experienced it as a… not a war, but a hostile environment*.*” (RCC)*

According to this interviewee, competition increased research individualism and dissolved the value of the scientific community.

Not too far from competition, **hierarchy** was another problem mentioned by some interviewees. In discussing with research students and technicians, we understood that hierarchies were inherent to academia and that they deeply influenced interactions, openness, and integrity. For example, both technicians and PhD students mentioned that they would find it very difficult to openly criticize the conclusion or dubious behaviour of the principal investigator of a laboratory. Most technicians mentioned that they would not dare to flag mistakes and errors because they felt that principal investigators were “*much smarter*” then them. PhD students said that they would refrain from disagreeing with a supervisor’s inadequate practice (i.e., we described a case of gifting authorship to a colleague who was not involved in the project) because they were worried that the supervisor would “*make it hard*” on them later on, and might even not allow them to graduate. Researchers, on the other hand, criticized issues linked with inequities in statute and reputation between researchers, saying that because of hierarchy in career achievements, “*the big will <become> bigger and the small will never have a chance*.”

Another relational component raised as a problem from the scientific culture was the issue of **unfair advantages from the research network**. Networking is an inherent part of science and was mentioned many times as an essential factor for success (34). Many researcher and students provided examples in which networking could help them get ahead, and some mentioned that there was comfort in knowing that good relations could bring ‘*favours*’ in case of need.

> *“Once you are in the network, you can also rely upon them and say ‘please do me a favour because I did a favour to you*.*” (Researcher)*
>
> *“…we were going for [a high impact journal], and then we were not writing the paper we were spending all our time trying to get the editor and the reviewers that they knew that the paper was coming so that once the paper was there all these people were involved and engaged and then it was either accepted or not. So it really depends on who you know and who you don’t know. And that’s why sometimes I start behaving like that… ‘Oh I would like to have a paper in one year in that journal’ and I start writing them and seeing them at conferences (laughs) ‘Hi, yes I’m thinking about submitting a paper what do you think about the idea’ and it really really helps. So it’s really not as unbiased as you would like it to be*.*” (Post-Doc)*

Nonetheless, most expressed discomfort, frustration, or loss of confidence from the advantages that research networks played on peer review for funding and publication.

> *“I feel like I have less and less confidence in publishing with the fact that ‘who is going to be the reviewer?’ ‘Is he biased?’ ‘Is it the journal?’ It’s like some politics that you I don’t always believe that the best results are published in the best journals*.*” (Researcher)*
>
> *“…it’s just the people that have the money that get the money. Because they’re all in the commissions or they have co-workers or close collaborators that are in the commissions, and they just give each other money all the time*.*” (RCC)*

Refuting these ideas, funders and editors sustained that their review process was organised to minimise conflicting interests. One editor mentioned that believing that good relationship with editors would help manuscript acceptance was “Wishful thinking”. She explained that strict policies against conflicting interests and the weight of external peer-reviewers in the decision cancelled what good networking could have created. To support her claim, she explained that she rejected the manuscript of her best friend not long ago. The discrepancy between the perspectives of researchers and the perspective of funders and editors makes this issue of unfair advantages difficult to resolve.

###### Punitive, not preventive

Added to the above problems, the worry that the scientific culture **focused on punishment rather than prevention** was raised by a few interviewees. Even though this issue was only mentioned by a few interviewees, their perspectives raised questions which are scarcely addressed in the current integrity discourse. A policy maker or influencer worried about the lack of a second chance for researchers convicted of misconduct. He sustained that once misconduct is proven, universities generally ban or shame the convicted researcher without offering any later contact, support, or chance for retaliation. He stated that “*This kind of unwillingness of the research system to forgive, not to forget, to forgive, really troubles me*.” This interviewee supported that in some cases, institutions would benefit from rehabilitating deviant researchers and involving them in integrity training later on. He believed that this would lead to higher relevance of integrity training, and would avoid that researchers who committed misconduct simply move on to a new university without any kind of follow up or notice — an issue that often happens in Europe where misconduct cases are not always disclosed publicly.

### A general resistance to change

In the final portion of our interview, we asked researchers ‘*who* they believe was responsible for promoting integrity’. Although selected actor-specific responsibilities were mentioned, we quickly realised that integrity was generally seen as a shared responsibility in which all actor have a role to play.

> *“So I think that it’s a broad ecosystem, and everyone has a role to play in that*.*” (EP)*
>
> *“I’m not going to say one person. I think it’s an extremely complicated theme, and extremely complicated idea, concept… So you cannot focus on one person. You need to target a lot of people*.*” (RCC)*
>
> *“Everybody [laugh]. Everybody has their share of responsibility, of course” (PMI)*

This sharing of responsibilities, however, appeared to downplay individual responsibilities and to trigger a shared feeling of **helplessness**. For example, researchers believed that, to survive in the current system, they had to play by the rules of the game, even if they disagreed with such rules. Institutions felt powerless on their own, and a former-researchers even believed that it was unrealistic to believe in any drastic improvements.

> *“Everyone is behaving like this. Everyone is saying ‘Let’s go for the safe road because this is how it is otherwise I will never get funding’, so…” (Researcher)*
>
> *“One institution cannot change that*.*” (RIL)*
>
> *“I don’t think we can expect, realistically speaking — but it’s cynical maybe — we can expect the great world change. It couldn’t change. You can try to make the ships sail a bit more in another direction but you cannot turn it. Therefore it’s too deep. The idea is… The views on what science is and how people work is too deep. It might be cynical if I’m saying it now*.*” (RCC)*

Lacking the empowerment or hope to take action, interviewees tended to transfer the root of the problems from one to another, creating a circle of blame which fostered frustration and distrust between actor groups. For instance researchers had to cut corners because universities pressure them to publish; universities had to push researchers to publish because policy makers distribute funding to universities based on publication outputs; policy makers had to distribute funding based on publications because society wants a return on its investment, etc. In other words, each actor appears to use the failures of higher actor groups to justify its personal inability to endorse best practices. But the complex interplay between actors also led to smaller circular criticism. For example, researchers criticized funders for evaluating them on quantity rather than on quality. But funders explained that even when they have policies in place to ignore quantity, peer-reviewers — who are themselves researchers — tended to cling to old quantitative metrics. Similarly, universities criticized that journals looked for hype rather than quality, but journals believed that the real problem was that universities used journals to evaluate researchers, not the decisions that journals take on what they choose to publish. Given that science is built around a community where all actors share the common goal of advancing knowledge, internal distrust and lost hopes for true change are necessarily a worry for the future.

### Short summary of findings

Our investigation of the problems that affect science and threaten integrity reveals a number of ideas on what needs to change in science. By involving different research actor in our analysis, we were able to discern the perspectives of different actors and to identify conflicting views, both between and within actor groups.

When discussing misconduct, interviewees sustained that misconduct was far from black and white. Indeed, the core reasons for condemning misconduct seemed to differ between individuals and actor groups. We also noticed that the jargon which is normally used to discuss misconduct and integrity was not common to all research actors. Finally, although ‘excessive pressure’ was the factor that was most often mentioned as causing misconduct, many sustained that the responsibility of misconduct ultimately resides in the researcher and that pressures cannot become excuses for bad practices.

We did not limit the discussion to strict misconduct. In fact, most interviewees were unfamiliar with genuine misconduct and were thus much more inclined to discuss the general problems which may deter research quality and integrity. In describing such problems, interviewees appeared to point to three general categories: Issues related to (i) personalities and attitudes, to (ii) awareness, and to (iii) the research climate. Issues related to personalities and attitudes were mentioned as potential targets for employers to consider, but were also admitted to be rather immutable. Issues linked to awareness generally discussed inadequate mentorship of research students and insufficient support on how researchers should meet integrity guidance. Finally, issues linked to the research climate highlighted problems which resulted from existing research environments and research cultures. The precariousness and scarcity of research careers, especially problematic for young researchers, were thought to be a major issue which aggravated the threatening impact of pressures and perverse incentives. Overspecialisation, expectations of versatility, and the lack of collaboration also came into play as constraining the time available for research, further intensifying the pressures on researchers and reducing the possibility for control and monitoring. Deeper in the cultures attached to research, the care and support given to researchers was also noted as being limited. Researchers were expected to participate in science out of passion, and thus to devote themselves without expecting personal benefits, but this perspective impacted the well-being and satisfaction of researchers. A general culture of profit, intolerance for failure, and expectations of extraordinary results added up to fuel a culture of ‘publish-or-perish’. The overwhelming pressure to publish further seemed to shape the relationships that researchers have with one another. Competitiveness, hierarchy, and alliances were described and believed to influence how research was planned, performed, and reported.

Finally, when asked about responsibilities for change, interviewees revealed a shared feeling of helplessness towards current problems. They felt that issues were caused by inadequate decisions of different actors, and thus felt frustrated and lost their trust in other actor groups.

## DISCUSSION

The present paper reveals a rich account of various stakeholders’ perspectives on misconduct and other problems of research and complements our associated findings on definitions and attribution of success in science (34). While it is technically impossible to integrate all diverse and sometimes inconsistent responses in a well-structured discussion, we would like to highlight three main findings from our paper series which provide insights for the next steps towards better science. Specifically, we will show how our findings align with the pressing need to i) revisit research assessment, ii) tackle inadequate climates and empower researchers, and iii) foster inter-actor dialogue.

First, the revision of research assessment needs to become central to the integrity discourse. Our respondents clearly indicate that definitions and assessments of success in science are not innocent, and that they impact research practices (34). While we understand the strong emphasis on metrics from a pragmatic point of view, in practice, our participants considered reductionistic metrics as imprecise, disruptive, and at the very heart of most problems afflicting science. Without discrediting excellent science that yields remarkable metrics, we must recognise that excellent science does not necessarily translate into such metrics and that current output metrics provide, at best, a reductionist picture of the qualities and merits of researchers. The lack of consideration for important research processes such as openness, transparency, and collaboration may dissuade researchers from investing in such practices which are ultimately essential for the quality and the integrity of science. In current climates, researchers who commit to good science regardless of short term impactful outputs may place their very existence as a scientist (i.e. their scientific career) at risk. Wide-spread expectations of extraordinary results further add to the problem, not only by suggesting that extraordinary science should be the norm — a paradox in itself — but also by devaluing negative findings and small-steps-science, both of which are key to advancing knowledge. And yet, current assessments were also said to ignore — even inhibit — high risk innovation, originality, and diversity. Considering all this, it is obvious that research assessment must be addressed. A number of recent initiatives, such as the Declaration on Research Assessment (DORA; 35), the Leiden Manifesto (36), the Metric Tide (37), or the Hong Kong Principles for Assessing Researchers (38), and numerous scientific editorials and public fora (e.g., 39, 40-42) are important pioneers in exposing the challenges of current assessments. Our findings echo these challenges and further link the problems to research integrity, thereby reinstating that research assessment must become central to the discourse on research integrity.

Second, our findings suggest that approaches to foster integrity should focalise on changing research climates rather than on individual behaviours. Our respondents sustained that research climates play a crucial role on research practices and integrity, a finding that is corroborated in most research on research integrity (33). Nonetheless, the majority of approaches aiming to tackle misconduct capitalise on the knowledge and awareness of researchers (33). Generally through training or codes of conduct, these approaches aim to “discourage scientists from behaving badly”. This person-centred perspective has profound implications on the way we perceptualize integrity. Not only does it ignore the dissonance between what researchers know they should do (i.e., integrity) and what helps them survive in their career (i.e., success), as described above, but it also transfers the burden of integrity on researchers — especially young researchers who are the main target of integrity training. In light of the high pressures, high demands, and lack of support that already afflict young researchers, it seems obvious that approaches to foster integrity need to better consider the climate in which researchers operate, the pressures it exerts, and the conflicts it entails. In this regard, training and education might need to shift their focus from *compliance* (i.e. what not to do) to *empowerment* (i.e., how to do great science) *and resilience* (i.e., how not to give in to cultural pressures which push you to alter your preferred approaches to scientific excellence). Training should aim to equip researchers to find ways to promote good science without jeopardizing their personal success as well as cultivate resilience and skills to allow smoother migrations from research to alternative careers. Likewise, support to and consideration of the person behind the research — something that has also been found missing in past works (e.g., 43) — should be prioritized. Ultimately, if young researchers who are adamant to good science are empowered and resilient to the current issues of academia, they will have a greater chance of surviving the precarious career cycle, of becoming activists for good science, and of shifting the tenacious cultures that currently disrupt integrity.

Finally, our findings reinforce the need for inter-actor dialogue in discussions on research integrity. When describing success in science, we sustained that a comprehensive inter-actor dialogue is needed to combine different meanings and expectations of scientific success (34). Similarly, when discussing problems of science with multiple actors, we understood not only that perspectives differ from one actor to the next, but that the lack of inter-actor discussion leads to a circle of blame in which no one feels able to tackle the problems. Even though actors depend on one another, the opportunities to discuss and share decisions between them are limited, especially for researchers. This segregation leads to misunderstandings, false beliefs, and missed opportunity for joint actions. As researchers, we were ourselves surprised to realise that pressures also affect institutions, funders, editors, and policy makers. We thus believe that the best way forward is to create fora for participatory decisions on topics of success, assessment, climates, and integrity. Prioritising opportunities for inter-actor dialogue and actively seeking the voice of overlooked actors will help reduce victimisation and blame and promote well-considered joint action.

## CONCLUSIONS

Our findings shed light on the complex interplay between success and research integrity. Involving not only researchers, but a wide range of actors who hold different roles in science, we show that there is great tension between what researchers should do to advance science, and what they must do to be successful. This finding resonates with debates that have been taking place in the past few years. But despite heated discussion, initiating changes in research assessments takes time, effort, and broad coordination. Our findings extrapolate a few action points which might help coordinate such changes. First, assessments of success must naturally be tackled and must become central to the integrity debate. Second, approaches to promote better science should be reassessed: not only should they consider the impact of the climate on research practices, but approaches which focus on researchers should also redefine their objective to empower and support researchers rather than to capitalize on their compliance. Finally, inter-actor dialogues and shared decision making are crucial to building joint objectives for change. Such dialogues should actively seek the voices of parties which are forgotten from the current discourse, and should genuinely aim to construct a collective understanding of the problem so that actors can join forces for change.

## Acknowledgements

The authors wish to thank Raymond De Vries, who substantially contributed to the Conceptualization, Methodology, Resources, and Validation of the present project. The authors also wish to thank Melissa S. Anderson and Brian C. Martinson and Raymond De Vries for sharing their focus group guides which constituted the foundation of ours (Resources). We also wish to thank Ines Steffens, Inge Thijs, and Igna Rutten who were essential in helping us organise focus groups and recruit participants (Resources).Finally, and most importantly, we want to thank all those who participated in our interviews and focus groups. We know that we forced ourselves in the very busy schedules of many a participant, and we are sincerely grateful for the time, efforts, and precious thoughts that participants generously shared with us.

## LIST OF ABBREVIATIONS

BOF: Bijzonder Onderzoeksfonds [special research funds]
EP: Editor(s) or publisher(s)
FA: Funding Agency(s)
LT: Laboratory technician(s)
PMI: Policy Maker(s) or influencer(s)
PostDoc: Post-Doctoral Researcher(s)
QRP: Questionable research practices
RCC: Researcher(s) who changed career
RIL: Research institution leader(s)
RIN: Research integrity network member(s)
RIO: Research integrity office member(s)

## DECLARATIONS

### Ethics approval and consent to participate

The project was approved by the Medical Ethics Committee of the Faculty of Medicine and Life Sciences of Hasselt University (protocol number CME2016/679), and all participants provided written consent for participation.

### Consent for publication

Participants provided written consent for participation and for publication and dissemination of the findings from this project.

### Availability of data and material

#### Data

Given the sensibility and risk of identification of qualitative data, we are unable to share full transcripts. In the manuscript however, we attempted to be as transparent as possible by providing numerous quotes in the text and in tables to exemplify and support our claims.

#### Materials

Our focus group and interview guides, as well as the consent forms and information sheet we shared with participants are available on the registration of this project (44).

### Authors’ contributions

Authorship contributions: NAB contributed to the Conceptualization, Project administration, Methodology, Resources, Investigation, Data curation, Formal Analysis, Visualization, Validation, Writing – original draft, and Writing – review & editing. WP contributed to the Conceptualization, Funding acquisition, Methodology, Resources, Validation, Supervision, and Writing – review & editing, both intermediate and final versions.

### Authors’ information

**NAB** is a PhD student supervised by WP at the faculty of Health and Life Sciences in Hasselt University, Belgium. Her research interests are predominantly in research integrity and publication ethics, especially targeting research assessments and the way in which scientific systems influence the integrity and the quality of science.

**WP** is associate professor at the faculty of Health and Life Sciences in Hasselt University, Belgium. As a medical ethicist with expertise in research ethics, he has increasingly devoted attention to education and research in the field of research integrity, with a specific interest in research culture and the definition of success in science. He is supervisor of NAB.

## Footnotes

1. In Flanders, a portion of the federal funding for research institutions is distributed using a special key called the BOF (bijzondere onderzoeksfonds [special research funds]) key (45). BOF funding is structural funding from the Flemish government which is distributed between the universities and research institutions in Flanders. Similar funding distribution systems exist in other countries around the world (46, 47). Once they receive BOF funding, Flemish universities are free to choose how they allocate the funds within their institution. The BOF-key is the calculation which determines the share of governmental research funding that each Flemish University will receive. Until 2019, the BOF-key was based on five indicators: master’s degrees awarded, defended doctorates, gender diversity, publications, and citations, with the latter two composing over 40% of the final score for distribution (47).

2. Since the interviews have been conducted, a new version of the BOF key has been developed and released (mid 2019). Nonetheless, a large proportion of the resource allocation distributed through the BOF-Key still depends on output metrics and publications with an impact factor.

3. Working beyond the 9–5 schedule was even seen as a factor for success (34).

