## Appendix 1 for "Rethinking success, integrity, and culture in research (Part 2) — A multi-actor qualitative study on problems of science"

### APPENDIX 8

#### SELECT QUOTES TO ILLUSTRATE THE PROBLEMS BEHIND MISCONDUCT AND 'BAD PRACTICES

##### Changing the course of science

"I suspect that those tiny [fraudulent] publications, they will get falsified, but nobody puts too much attention to that anyway, **they are not moving the real direction of science, you know, this is such a small study somewhere, it doesn't really matter that much** if it turns out to be incorrect. [...] Cutting corners sometimes is not changing the course of science. [...] So it can happen to everybody that you cut corners. It's just, is that a scientific corner or is that an esthetic thing? But everybody will be tempted at some stage to do that so let's just hope that the majority of people will have the scientific integrity not to do that. But... (sighs) I think, certainly with PhD students I only hear transiently... But I don't think you can get a full article based on one little corner that has been cut. I think you really need to cross that line, and then go up and not do that again. I think that's where things go wrong when it becomes a standard, where you've made it once and realised it doesn't make a difference..." (RIL)

##### Changing conclusions

"In my lab if I look, the only misconduct I've picked up was just stupidity. PhD students who scanned a little too short and had to go back to the scanner and thought "I could just copy-paste the bottom bit because there's nothing on it anyway". **That's real misconduct, but at the same time, that's not scientific fraud. Well it was, it is scientific fraud, but he was not changing a conclusion**, he was just too lazy to scan a really nice experiment [...] What I consider cheating is that you leave out the data that don't suit your model. Or you make up data to get your model correctly. That is what I call cheating."(RIL)

"It's difficult to prove intention, and **for us that's not that important. If it's actually a deception, doesn't matter if it's intentional or not**. Then we need to correct the literature if it's published. So you know, again I think a lot of the misconduct investigartion that institutions do, they put a lot of emphasis on the intentional bit because that's part of the employer status and so on. Whereas we as journals are not that interested in that part. **We're really interested in 'Is this research trustworthy or not?**" (EP)

"Intent is something that editors are not in a position to properly evaluate. [...] And this is where they have to rely on institutions to determine, to really establish and ascertain whether there is misconduct or not. Where the responsibility of the editor is in **correcting the scientific record**. And that, it doesn't matter whether it's misconduct or not in a way.[...] **if something is wrong**, and you're unsure as an editor **whether it's misconduct or not, it doesn't matter**. It also needs to be corrected in the scientific record." (EP)

"I think usually that's something that is important in evaluating those cases. It's really, does the act, the problem that you have identified, **does that actually lead to a different... To change the nature of the conclusion?** And really make the data say something different than what it says? And so… You know that tends to be misconduct." (EP)

##### Poor quality of findings

"You always have to take into account the rules, the procedures of good research.[...] If you want to talk about society, your reference group must be big enough so it can be a reference, a real reference to the society. So if you don't put up the research in advance in a good way, then **it's also sloppy or bad research. Because all the results that come out of this, even if they're positive, they won't be representative for the reality.** And that can lead to — if it's research in medicine—that can be very dramatic. So that's sloppy research." (PMI)

##### Misuse of research money

"About copying deliverables for a different research grant "Yeah, well, for us it's fraud directly. Because **you do it in order to win money**." (PMI)

| Intentional |
|---|
| "For me a bad scientist is someone who actually intentionally knows that what he or she is doing is wrong and might inflect the public opinion upon a publication. […] So for me, being a bad scientist **actually means intentionally being bad**. You can perform a research without being aware [of the rules] [...] That doesn't make you a bad scientist or a bad researcher. " (RIO)<br><br>"If someone warned you that this is not okay, that it is bad science, and **you still continue**, then, yeah, it's also not okay." (RIO)<br><br>"There is big misconduct or minor misconduct, if we can put it like that. It's like when you **consciously know** you are really doing some change in your results to make them look beautiful and then get this publication in nature, for whatever reason, or when you are just tweaking here and there and the supervisors is telling you everybody does that, so you are able to do that [laughs]. There's **different degrees of seriousness** also in here." (PMI)<br><br>"Well if it's **willingly** then it's a… It's a border you don't cross. (PMI) |
| **Moral mismatch** |
| "[Sometimes researchers say] 'yeah but it didn't change the main results of my article, so what's the problem?' [...] OK **if the results are being the same, that's not the issue actually**, it should be the process also. And at that point you see that there is this **moral mismatch**." (RIO) |
