## Appendix 2 for "Rethinking success, integrity, and culture in research (Part 2) — A multi-actor qualitative study on problems of science"

### RANGE OF RESPONSES TO THE QUESTION ON '*WHY* MISCONDUCT AND QUESTIONABLE RESEARCH PRACTICES HAPPEN'

The number of quotes and interview should be considered with caution since they depend on the capture and the coding, yet we found interesting to show it as a rough estimate of the coverage of select themes.

| Topic | # of quotes | # of interviews | Sample quote | Actor |
|---|---|---|---|---|
| **Pressure** | 29 | 22 | ...pressure for career, reputation is playing a role, competition is playing a role... that's all external. | RIL |
| | | | I can imagine that if you're in a situation where you're forced to have a certain outcome and your reputation depends on it – and I find it very shameful that there are, or there might be situations like that – that as a researcher you try to bend the truth in your favour. [...] I do believe that there can be situations where stress might force you to go into a direction that you wouldn't have walked in normally. | RIO |
| | | | The pressure is huge. You know, basically your career depends on it. And so when you are in a system in which basically your next paycheck, your next grant will be dependent on the results, I've never thought about it, but I can imagine that you will have some people think 'Well… Why don't just, you know, make it up?'. | PE |
| | | | They need their numbers of publications. [...] Pressure, yeah. | PMI |
| | | | …the more mundane reason I think has a lot to do with time and publication pressure and the pressure for funds, fundraising and things like that. Which puts an enormous pressure on people to produce results, to publish results… | RF |
| **Ego and personal morals** | 23 | 12 | Internal it's idolness, the personal 'I want to be a big researcher', you always should be modest as a researcher I think. So the lack of modesty is the internal factor that makes people just improving their data a little bit., just adding something here and there, deleting something here and there… | RIL |
| | | | I think that there is the egos, and the egos in science is still underestimated I think. yeah... [...] if you look at it when professors are doing it [i.e., committing misconduct], then it has a lot more to do with status, image, ego, trying to score, get off easy, yeah... I see that a lot more than it is of the pressure issue. [...] it's about image and scoring. | RIO |
| | | | But the ones that really matter, that make it to the press, those are very often the leading universities and I think there the perverse incentives are big egos, big prizes, top publications... There, you know, if you manipulate your data, you become a big hero. And I think that's at the level of the promoter, and that's probably the biggest problem of them all. | RIL |
| | | | You know that when you start fabricating papers to have like two papers a year in Nature, you don't do this for policy reasons. [...] You do this because you want to be kind of a king or a god in your discipline, and that's… well maybe narcistic, maybe psychopathic type of behaviors. | PMI |
| | | | I think that depends on the person. (laughs)… | PMI |
| **Normalisation of smaller misbehaviours** | 11 | 10 | Because they start with QRP, and it gets more and more and more, and then they cannot admit it anymore without seeing the consequences so they make the choice to do even more terrible fraud to cover up the rest. | RIL |
| | | | And also the fact that it's a slippery slope, in fact. And that when you start being a little sloppy about certain things, you can actually very easily drift into something that is much worse. | EP |
| | | | I think if you get away with small enfringements then you get bolder and bolder each time and, you know that happens. | EP |
| | | | And when from the moment you do it, you do it again, and when you think your colleague does something you will do it again etc. | RCC |

| | | | | |
|---|---|---|---|---|
| **Perverse incentives** | 8 | 7 | I think we have a lot of perverse incentives. […] But, when you are a student, and your four years are up, your promoter does not have money to pay you any longer, and there's nothing you have to publish, and there's one excel spreadsheet if you change a few numbers, will give you a publication, I think it's very tempting to do that. | RIL |
| | | | if you actually do experiment with small numbers of animals, you're going to have a much larger effect. […] These are the kinds of perverse incentives that we have in the system at the moment. | EP |
| **Lack of awareness** | 7 | 5 | But people come willingly and tell us "It's the first time that I hear about the codes. It's the first time that I hear about those things". | PMI |
| | | | I don't believe that there are researchers who intentionally perform bad research, I do believe that there might be some researchers who are not aware of what the better practice or the best practice might be. And if they are informed about the better and the best practices in specific research and they follow these practices and they adjust their work methods, that's not a bad scientist. | RIO |
| | | | What I think is important though, is that, again in my generation, we were not made aware enough of that. A bit like gender bias; until you're explained what gender bias is, until you're explained what research integrity is, and what misconduct is, I think you will have more flexibility towards it because you just don't know where the red line is. Once you explain to people where the red line is, they will know when they cross it. And I guess… I think, they will cross it less likely. | EP |
| **Lack of control (low risk high gain)** | 5 | 5 | "People don't see the severity of [mibehaviors like gift authorship]. So then the gain is much bigger than the risk." | RIL |
| | | | ...you know that indeed if you do that, you will get a top publication which is very beneficial for your career. | RIL |
| | | | While big industrial laboratories have standard operating procedures that are very expensive and standardised when it's not easy, in research institutions, we are missing them. [...] Sometimes this is missing. Right? So that can lead to sloppy research. | PMI |
| **Unrealistic demands** | 4 | 2 | But at the end of the day I think the motivations are fairly similar for sloppy research and misconduct. And I think, in my view, it's related to the first question we discussed, which was the question of incentives, and research assessments. [...] Success is measured by publishing in very selective journals that are looking for very important ground-breaking claims. You are incentivised to find these ground-breaking claims… And so you are incentivised to really get something that is extraordinary, and ground-breaking. And let's face it, all the research in biomedical research, is not ground-breaking and extraordinary. Most of it is not. | PE |
| | | | I think there are too many PhD students who are really forced into working 24/7, which definitely cannot be what it should be either… | RIL |
| **Lack of openness to failure and negative [illegible]** | 3 | 2 | If a researcher does every step of the research process correctly, and after ten years, he only had negative results, will the management of the institution still fund him for the next five years without guarantee that there will be a positive result? Or will they say after ten years 'Now it's enough you stop. You can go and find another job.'? In the latter case if that's the case, then I think every researcher, theoretically, every researcher can be changing its data. | PMI |
| | | | And if we do not tolerate failure, and use failure as a motor to drive you to success, you foster misconduct and sloppy research. Most of my researchers would be very upset, but that's how it is! | PMI |
| **Overspeci alisation** | 2 | 2 | ...there's also this hyper-specialisation where people always go further and further and become more sophisticated within a paradigm, without even questioning the paradigm anymore. | RF |
| | | | In all research misconduct that have been analysed, there are usually three that are present in all. The researcher always knew better. He was under pressure. And he was in a research area that was very difficult to replicate. | PMI |
| **Cultural background** | 2 | 2 | These were two [foreign] post docs who faked some western blot data I think and I think that this was a very ambitious lab with a lot of pressure. [I know people from this nationality] so I dare to say that [they] have a slightly different opinion about rules. So they're more relaxed than [western Europeans] for example, and well... I think they didn't think it's that serious. | RIL |
| | | | ...he/she doesn't mind plagiarising as long as he/she doesn't get caught. So yeah... That also had to do with his/her nationality. [...] I think [some cultures] have this mentality that it's almost, you honor somebody by plagiarising them. And they just want to get their diploma so they can do a post doc in America. And he already had a post doc lined up. So he was really annoyed that he now had to postpone his post doc by a couple of months... | RIL |
